# Diffusion-limited cytokine signaling in T cell populations

**DOI:** 10.1101/2024.01.18.576190

**Authors:** Patrick Brunner, Lukas Kiwitz, Lisa Li, Kevin Thurley

## Abstract

Effective immune-cell responses depend on collective decision-making mediated by diffusible intercellular signaling proteins called cytokines. Here, we designed a spatio-temporal modeling framework and a precise finite-element simulation setup, to systematically investigate the origin and consequences of spatially inhomogeneous cytokine distributions in lymphoid tissues. We found that such inhomogeneities are critical for effective paracrine signaling, and they do not arise by diffusion and uptake alone, but rather depend on properties of the cell population such as an all-or-none behavior of cytokine secreting cells. Furthermore, we assessed the regulatory properties of negative and positive feedback in combination with diffusion-limited signaling dynamics, and we derived statistical quantities to characterize the spatio-temporal signaling landscape in the context of specific tissue architectures. Overall, our simulations highlight the complex spatiotemporal dynamics imposed by cell-cell signaling with diffusible ligands, which entails a large potential for fine-tuned biological control especially if combined with feedback mechanisms.

## Introduction

Interactions between immune cells play a fundamental role in the mammalian defense against pathogens. Specifically, the fine-tuned decision-making processes in the adaptive immune response comprise cell-cell communication amongst antigen-presenting cells (APC) and T and B lymphocytes, employing surface-mediated signaling as well as diffusible ligands called cytokines^1,2^. Remarkably, different cytokine species may share important parts of the signaling machinery, including subunits of the high-affinity cytokine receptor as well as downstream signaling mediators, and still evoke different biological functions and regulatory properties. For instance, the cytokines Interleukin(IL)-2 and IL-7 share the common gamma-chain of their receptors, and they both utilize STAT5 as major signaling mediator^3^. While IL-2 plays an essential role in the regulation of T cell activation as well as clonal expansion, IL-7 controls the homeostatic T cell population size. Furthermore, IL-2 signaling causes an increased expression of the high-affinity IL-2 receptor (IL-2R) on target cells^4^, while IL-7 signaling causes down-regulation of IL-7 receptor (IL-7R) expression in T cells^5^. Due to its promoting effect on T cells, low-dose IL-2 therapy has been successfully employed in cancer immunotherapy and for autoimmune diseases, while IL-7 is utilized in the treatment of infectious diseases^6–9^.

In the case of paracrine cytokine signaling, the cytokine sources and sinks are often separated^10^, which may result in a spatially uneven cytokine concentration due to consequences of diffusive cytokine transport^11^. Indeed, previous model simulations have predicted spatial inhomogeneities in cytokine concentration spanning several orders of magnitude, within a physiological parameter regime^12–14^. Those results have been supported by experimental findings of notable and tunable inhomogeneities of the cytokine concentration in secondary lymphoid organs, which regulate the formation of local cytokine micro-environments^15–18^. Nevertheless, the measured fast diffusion coefficients for cytokines such as IL-2 may counteract the effects of localized cytokine secretion and uptake^17,19^, suggesting a subtle balance of several regulatory mechanisms controlling the spatial distribution and signaling range of cytokines. In fact, theoretical as well as experimental studies have indicated that spatial cytokine inhomogeneities can be fine-tuned by properties of the cell population^13,16,20^, but the range and effect size of such mechanisms remains unclear. Furthermore, it is not known how different regulatory mechanisms employed by different cytokines, such as positive vs. negative feedback on receptor expression in the case of IL-2 and IL-7, specifically modulate their spatio-temporal signaling properties and how that relates to biological functions.

Next to the emergence and fine-tuned control of spatial cytokine inhomogeneities, a prevailing question concerns the consequences of cytokine inhomogeneities for paracrine signaling efficacy and, in more general terms, for decision-making processes of immune-cell populations. Intuitively, spatial cytokine inhomogeneities may promote signaling efficacy especially in the range of low average cytokine concentrations, because locally enriched concentrations in small microenvironments may act to overcome the threshold for signal induction at least in those areas. However, immune-cell populations are subject to complex non-linear dynamics and constraints imposed by the detailed tissue architecture. All those system properties act together in shaping the spatio-temporal dynamics, and therefore have to be considered in a quantitative analysis of the spreading and efficacy of paracrine cytokine signals.

Here, we designed a precise yet flexible mathematical simulation platform based on the finite-element method, to systematically analyze the interplay of tunable regulatory properties of paracrine cytokine signal propagation. Our investigation revealed the number of cytokine secreting cells as the primary driver of inhomogeneities in the cytokine field. Furthermore, feedback mechanisms involving receptor expression for both IL-2 and IL-7 finely regulate the activation of cells around a cytokine source, which we quantified in the model simulations by developing specifically tailored summary statistics. Finally, as the model allows for a variety of cytokine dynamics and interacting cells, we explored the consequences of specific tissue architectures on cytokine distribution and signaling. Throughout those multiple levels of paracrine interaction, we found that the complex diffusion and uptake dynamics generate properties of signal propagation that are quite different from a well-mixed scenario ignoring spatial inhomogeneities that is studied in parallel.

## Results

### Spatially inhomogeneous cytokine concentrations arise generically and can induce potent and fine-tuned paracrine signals

Effective paracrine cytokine signaling requires concentrations exceeding a threshold for signal reception at the target cell^21^, and therefore is in conflict with the low measured values for the average concentration of many cytokine species^13,22^. Diffusion-limited signaling is a plausible mechanism for effective paracrine signaling even under conditions of low tissue-level cytokine concentrations, because locally amplified concentrations in the vicinity of cytokine-secreting cells may allow to exceed the signaling threshold on the surface of nearby responder cells. Nevertheless, such a mechanism requires a subtle balance of the rates of cytokine secretion, cytokine uptake and cytokine diffusion, supplemented by intracellular processes including signaling cascades and transcriptional regulation. To systematically assess the robustness and dynamic range of diffusion-limited cytokine signaling in the context of regulatory properties of a cell population, we designed a spatio-temporal simulation work-flow based on an accurate and efficient finite-element solver. Due to the large time-scale separation between diffusion (seconds), intracellular signaling (minutes) and processes that require transcriptional regulation (hours), we decided to employ a quasi-stationary state assumption to the reaction-diffusion problem throughout. Since we are interested in the systematic analysis of local cell-cell interactions, special consideration was put into a modular, scalable and parallelizable modeling environment (Figure S1A). For the chosen mesh configuration, computation time scaled linearly with mesh fidelity and system size (Figure S1B), and the simulation error was largely set by boundary effects (Figure S1C-E) and therefore decayed rapidly with system size.

To set the stage, we initially studied direct paracrine signaling activities in an otherwise stationary cell population, in terms of the well-studied model system of IL-2 secretion and uptake by T helper cells in a scenario with randomly assigned cytokine secreting and responder cells (Figure 1A). Model parameters were determined in line with experimentally measured quantities ^4,12,13,19,23–25^ (Table 1), to foster simulation results in the physiological parameter regime. Following previous work^16^, we accounted for saturation effects regarding cytokine binding and uptake by cytokine receptors using a Michaelis-Menten type of equation, which we found to be a direct consequence of a mechanistic model formulation (Supplementary Text, “Description of uptake dynamics”). In line with experimental data^14,25,26^, we assumed a discrete all-or-none type of IL-2 secretion in cytokine-secreting cells (the impact of this assumption is studied in detail below). We additionally considered intracellular signal transduction by means of a conceptual model, where the phosphorylation level of the signal transducer and activator of transcription (STAT) in responder cells indicates effective paracrine signaling.

**Figure 1:**
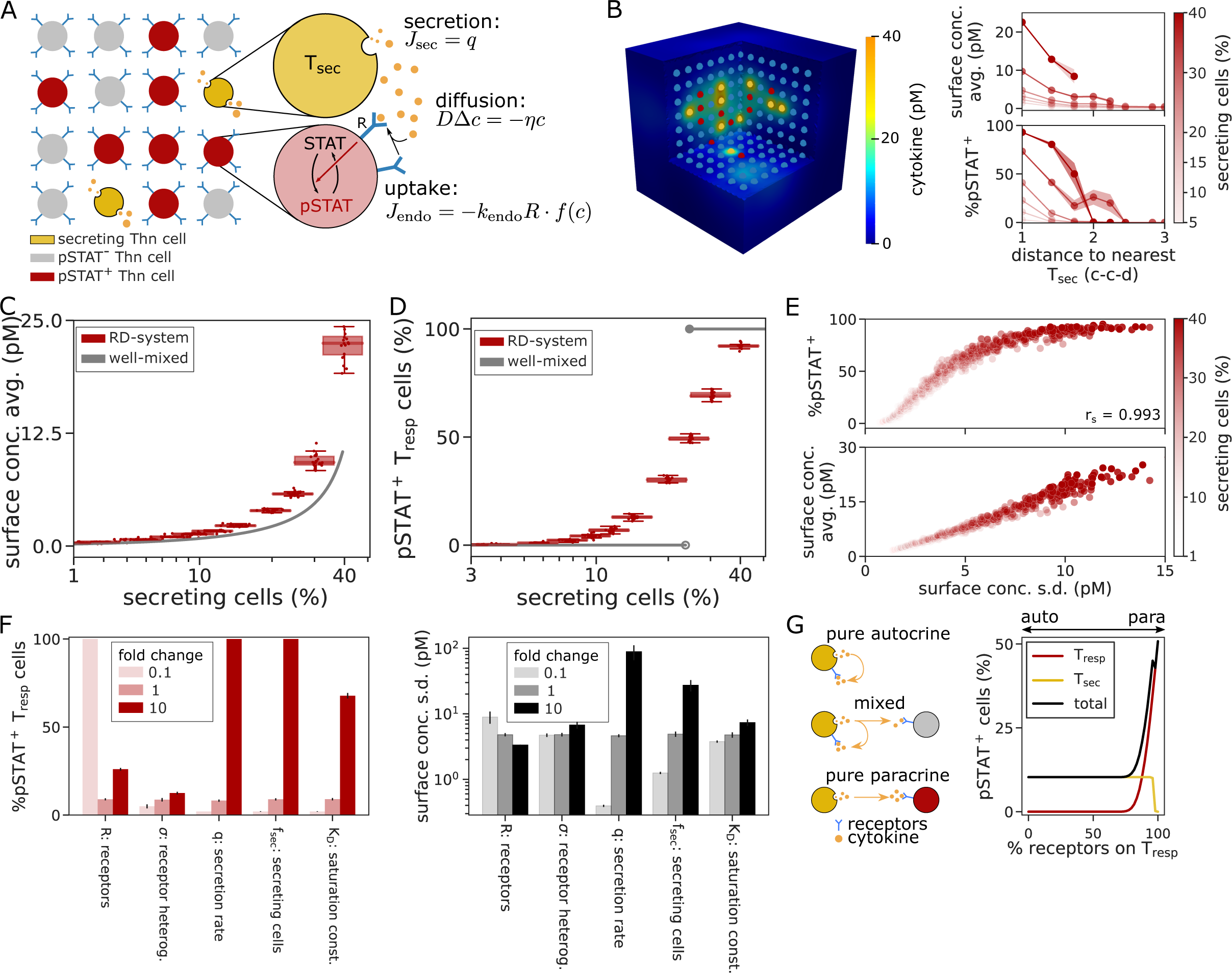
Spatial inhomogeneities induce stable paracrine signaling activity. (**A**) Model scheme. Cytokines are released by the secreting cells, and are subject to diffusion in the extracellular space and to binding to receptors on Thn cells. This binding and complex building is then translated into the phosphorylated STAT signal. (**B**) Typical model simulation using standard parameters (Table 1) and 5% secreting cells. Shown are the cytokine concentration field (left, see also legend in panel A) and the cytokine concentration and resulting pSTAT5 levels on responder cells (right). c-c-d: cell-to-cell distances. Maximum distance: 50% of average-distance between secreting cells. (**C-D**) Average cytokine concentration on the cell-surface across responding cells and percentage of pSTAT+ cells for varying fractions of cytokine secreting cells, in the RD-system and corresponding well-mixed scenario, as indicated. Cells with pSTAT > 0.5 are considered pSTAT+. (**E**) Correlation between the fraction of pSTAT+ cells and the average surface concentration as shown in panels C and D, with spatial inhomogeneity measured by the standard deviation of the cell-surface concentration across responder cells (spatial s.d.). Each dot represents a single simulation run. rs: Spearman’s rank correlation coefficient. (**F**) Sensitivity analysis with respect to the fraction of pSTAT+ cells and the spatial s.d. (see panels C-E). The x-axis indicates the parameter varied by a factor (fold-change) as indicated, all other parameters remain constant (cf. Table 1). (**G)** Scan from pure autocrine (all receptors on secreting cells) to pure paracrine (all receptors on responder cells) signaling as indicated by the cartoon. Shown is the percentage of pSTAT+ cells for responding cells (red), secreting cells (gold) or both (black). Lines with shaded regions (panels B and C) and errorbars (D and F) indicate averages and standard error of the mean (SEM) across 20 runs of the model simulation.

**Table 1:** Standard parameter values.

| Parameter | Value | Unit | Description | Source |
| --- | --- | --- | --- | --- |
| $N_{\text{cells}}$ | 1000 | | Amount of cells in the system | |
| $d_c$ | 20 | $\mu\text{m}$ | Shortest distance between cell centers | |
| $r_c$ | 10 | $\mu\text{m}$ | Cell radius | |
| $D$ | 10 | $\mu\text{m}^2/\text{s}$ | Diffusion coefficient | 19 |
| $I$ | $10^{-12} - 10^{-9}$ | M | Cytokine concentration range | 4 |
| $f_{\text{sec}}$ | | | Fraction of secreting cells | |
| $f_{\text{resp}}$ | | | Fraction of responding cells | |
| $k_{\text{on}}$ | $3.1 * 10^7$ | $1/(\text{M} * \text{s})$ | Cytokine-receptor association rate | 23 |
| $\eta$ | $2.8 * 10^{-5}$ | 1/s | Cytokine decay rate | 12 |
| $\sigma$ | 1 | | Receptor heterogeneity | 4 |
| **Saturation parameters** |  |  |  |  |
| $k_{\text{off}}$ | $2.3 * 10^{-4}$ | molecules/s | Receptor disassociation rate | 23 |
| $k_{\text{endo}}$ | $4.6 * 10^{-4}$ | molecules/s | IL-2*R complex internalization rate | 23 |
| $K_D$ | $7.423 * 10^{-12}$ | M | Concentration of half saturation | |
| **Receptor feedback** |  |  |  |  |
| $k_{\text{base}}$ | $1 * 10^{-3}$ | molecules/s | Minimum receptor production rate | |
| $v$ | 0.042 | molecules/s | Receptor decay rate | 13 |
| $K_m$ | 0.5 | | pSTAT5 value of half feedback | |
| $\gamma$ | 0.01 - 100 | | Feedback fold change | 4 |
| **pSTAT signaling** |  |  |  |  |
| $K_{\text{EC50}}$ | 860 | Molecules | Receptors of half EC50 response | 24 |
| $E_{\text{max}}$ | $125 * 10^{-12}$ | M | Maximum EC50 concentration | 4 |
| $E_{\text{min}}$ | 0 | M | Minimum EC50 concentration | 4 |
| $N_{\text{EC50}}$ | 0.55 – 1.5 | | EC50 hill exponent | 4 |
| $N_{\text{pSTAT5}}$ | 3 | | pSTAT5 hill exponent | 24 |
| **IL-2** |  |  |  |  |
| $q_{\text{sec}}$ | 10 | molecules/(cell * s) | Secreting cells IL-2 secretion rate | 25 |
| $R_{\text{sec}}$ | 100 | molecules/cell | Secretor cells receptor count | 4 |
| $R_{\text{resp}}$ | 1500 | molecules/cell | Responder cells receptor count | 4,24 |
| **IL-7** |  |  |  |  |
| $q_{\text{sec}}$ | 100 | molecules/(cell * s) | Secreting cells IL-7 secretion rate | |
| $R_{\text{sec}}$ | 0 | molecules/cell | Secretor cells receptor count | |
| $R_{\text{resp}}$ | $5 * 10^5$ | molecules/cell | Responder cells receptor count | |

Our modeling setup resulted in a high degree of spatial patterning due to cytokine concentration gradients between secreting and responding cells, which were also reflected in the downstream STAT signal (Figure 1B). As anticipated, our simulations revealed appreciable paracrine signaling efficacy due to increased local cytokine concentrations (Figure 1C and D). Quite remarkably, high paracrine signaling activity (up to 40% pSTAT+ cells) occurred in a regime where such signaling was undetectable in an identically parameterized ordinary differential equation (ODE) system (Figure 1C and D, “well-mixed”) that arises naturally from a fast-diffusion assumption (Supplementary Text, “Deriving the well-mixed model”). Moreover, diffusion-limited signaling generated an appreciable dynamic range with regard to regulation by the amount of cytokine secreting cells, while the system was limited to an all-or-none response in the well-mixed situation. Accordingly, paracrine signaling efficacy exhibited a strong correlation with an increase in spatial cytokine inhomogeneity as quantified by the concentration gradient (Figure S2A) or the spatial standard deviation (s.d.) (Figure 1E and S2B) of cytokine levels across responder cells.

To characterize our model system in more detail, we performed a systematic parameter sensitivity analysis with respect to standard parameter values (Table 1). That analysis revealed strong effects on signaling efficacy and cytokine inhomogeneity by the receptor number and the half-saturation constant for cytokine uptake KD in addition to the cytokine secretion rate and the fraction of secreting cells (Figure 1F, Figure S2C-D). Interestingly, the rates of diffusion and extracellular cytokine decay as well as the cell-to-cell distance have a minor effect on signaling efficacy and cytokine inhomogeneity. Hence, signaling amplification by diffusion-limited cytokine propagation is a generic mechanism that is robust to subtle changes in the spatial configuration of the system, but sensitive to properties that are under control of the participating immune-cell populations.

In line with that, experimental evidence^10^ and theoretical considerations^19^ suggest the ratio of receptors between secreting and responding cells to be a carefully controlled property that determines the mode of cytokine signaling in the range between purely autocrine and purely paracrine signaling (Figure 1G). In our simulations, we found that paracrine signaling is limited to situations with more than 75% of all cytokine receptors expressed on responder cells and a subsequent increase in spatial inhomogeneity (Figure 1G, Figure S2E and F). On the other hand, cytokine secreting cells require only a minimal amount of receptor expression to receive an appreciable cytokine signal. That discrepancy is in line with the requirement for careful control of paracrine inflammatory signals such as IL-2 in order to prevent a potentially lethal cytokine storm^27,28^, and may explain the previously observed^29^ down-regulation of the pSTAT5 signaling pathway in cytokine secreting cells.

Overall, we found that a diffusion-limited mode of cytokine signaling allows for effective paracrine signaling in a regime of low average cytokine concentrations, and contrasts with a well-mixed scenario in the same parameter regime where paracrine signals remain far below thresholds for onset of downstream signaling cascades.

### Fractional abundance of cytokine secreting cells as major source of spatial inhomogeneity

Having established the generic occurrence and tunability of diffusion-limited paracrine signaling amplification, we sought to investigate the contributions of individual system components to cytokine concentration inhomogeneity. In our simulation, a uniform cell population, where cytokine secretion and uptake were equally distributed across cells, generated a nearly homogeneous cytokine concentration field, despite localized secretion and uptake at the cell surfaces (Figure 2A). Hence, we expected that a tight localization of cytokine sources is critical for spatial cytokine inhomogeneities, and we further sought to analyze the contributions of a heterogeneous distribution of cytokine receptors and of saturation effects in cytokine uptake dynamics on responder cells (Figure 2A). To this end, and to test the impact of an all-or-none-behavior of cytokine-secreting cells (that is few cells secreting large amounts of cytokine), we designed a simulation setup in which the total amount of secreted cytokine molecules and of cytokine receptor expression remain constant under parameter variation. We found that increasing the number of secreting cells in that system resulted in a steep decay of concentration inhomogeneities, approaching the well-mixed scenario (Figure 2B, left panel). The rise in the average cytokine concentration for low amounts of cytokine secreting cells (<5%) could be attributed to saturated uptake rates at high local concentrations, as it disappeared in the corresponding system with linear uptake rate (Figure S3A), in contrast to the increase in spatial inhomogeneity which occurred also under linear uptake.

**Figure 2:**
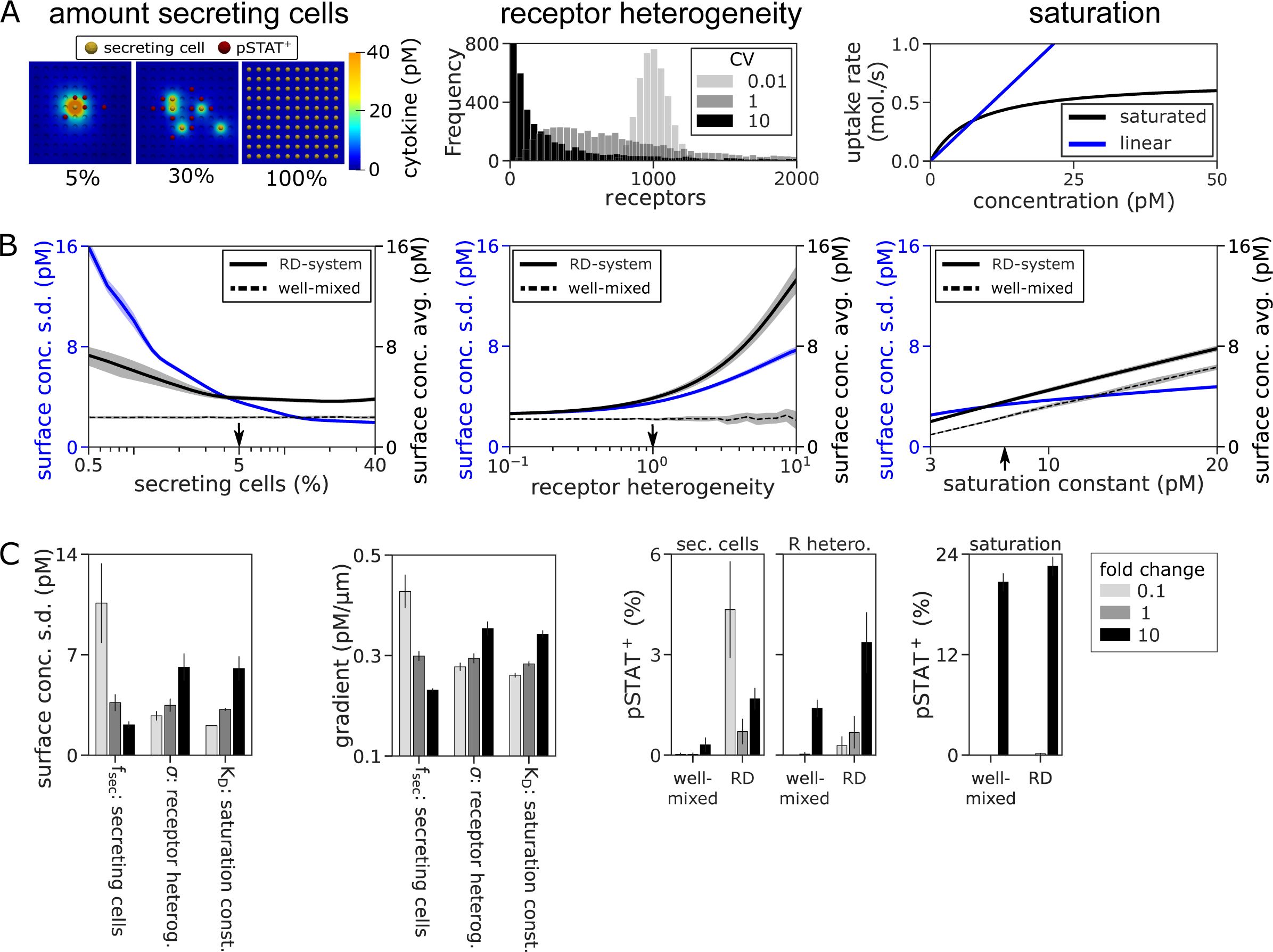
Systematic analysis reveals major drivers of cytokine inhomogeneity. (**A-B**) Analysis of cytokine inhomogeneity with respect to the fraction of cytokine secreting cells, receptor heterogeneity and saturation constant as indicated. In contrast to the analysis in Figure 1, simulations are designed in such a way that the total number of secreted cytokine molecules and the total number of cytokine receptors in the system are conserved through all simulations, see text for details. Shown is (A) a visualization and (B) a systematic scan for the parameter under study with respect to the spatial s.d. and spatial average (cf. Figure 1C), all other parameters are kept at standard values (Table 1). Vertical arrows indicate standard parameter values. (**C**) Sensitivity analysis of the parameters under study with respect to spatial s.d., gradient and fraction of pSTAT+ cells, analogous to Figure 1F. Lines with shaded regions (panel B) and arrow (C) indicate average and standard error of the mean (SEM) across 20 runs.

To account for cell-to-cell heterogeneity in cytokine receptor expression^4,24,30^, we considered expression values following a log-normal distribution at varying coefficients of variation, while keeping average expression levels constant (Figure 2A, middle panel). As anticipated, high levels of receptor heterogeneity induced cytokine inhomogeneities in the RD-system, but not in the well-mixed scenario (Figure 2B, middle panel). Interestingly, in the RD-system, the average cytokine concentration also showed a substantial increase with receptor heterogeneity, although the total amounts of cytokine secretion and uptake were kept constant so that cytokine concentrations remained unchanged in the well-mixed scenario. That seemingly paradoxical effect is independent of uptake-rate saturation (Figure S3B). It can be intuitively explained by a lower chance for high uptake capacities in the vicinity of cytokine secreting cells, which we could analytically reconcile by the help of a previously established^13^ approximate solution to the reaction-diffusion problem (Supplementary Text and Figure S3C).

Finally, we considered the effects of varying the saturation constant in our system. Of note, substantial differences regarding average cytokine concentration values between linear and saturated uptake functions occurred only in parameter ranges with very high secretion rates (Figure S3D), which are reached at low fractions of cytokine-secreting cells (cf. caption to Figure 2B). In line with that, increased values of the saturation constant KD induced only moderately higher values of the spatial inhomogeneity, and a nearly proportional change in concentration (Figure 2B, right panel), in both the RD and the well-mixed system. Across all three parameter values under study, we found that any increase in inhomogeneity was accompanied by a similar rise in activation (Figure 2C, Figure S3E and F). While that increased signaling activity can partially be attributed to the increased average cytokine concentration in all three cases, the change in cytokine concentration values is lowest and the increase in spatial inhomogeneity is the highest in the case of a decrease in the fraction of cytokine secreting cells.

Hence, we conclude that the highly localized mode of cytokine secretion in a situation with a small number of highly active secreting cells is the main driver of cytokine inhomogeneities, in turn increasing localized paracrine signaling efficacy.

### Dynamic feedback on receptor expression modulates spatial cytokine gradients

In addition to the mere exchange of paracrine signals studied so far, immune-cell populations have been found to exhibit feedback regulation directly on the level of cell-cell communication, in terms of cytokine receptor expression levels depending on the local cytokine environment^5,31–33^. Therefore, we proceeded to study the impact of such feedback mechanisms on the cytokine concentration field. Since modulation of receptor expression levels requires transcriptional regulation, considering such processes introduces a new time-scale on the order of hours to the system, giving rise to an intertwined combination of a quasi-stationary reaction-diffusion problem and a comparatively slow, dynamic modulation of cellular properties. In immune-cell compartments such as the lymph node, experiments have shown that several lymphocyte populations show high degrees of random and directed cell motility, in many cases achieving cell speeds on the same time-scale (hours). However, upon effective antigen stimulation, T cells remain bound to an antigen-presenting cell via the immunological synapse until they reach their full activation status by means of additional cytokine signaling^34,35^. That gives rise to an immobilized population of antigen-exposed responder cells, which is the focus of our study.

Positive and negative feedback on cytokine receptor expression are widespread properties of immune-cell populations, for which we designed a generic and reusable mathematical formulation using our established response-time modeling framework^36,37^ (Figure S4A). Here, we focused on two well-established examples, which are the IL-2/IL-2R system for positive feedback and the IL-7/IL-7R system for negative feedback (Figure 3A). In both cases, naive T helper cells act as responder cells.In the case of IL-2, cytokine secreting cells correspond to fully activated T cells under high antigen stimulation^7^, and in the contrasting scenario of IL-7 signaling, stromal cells take the role of secreting cells^5,38^.

**Figure 3:**
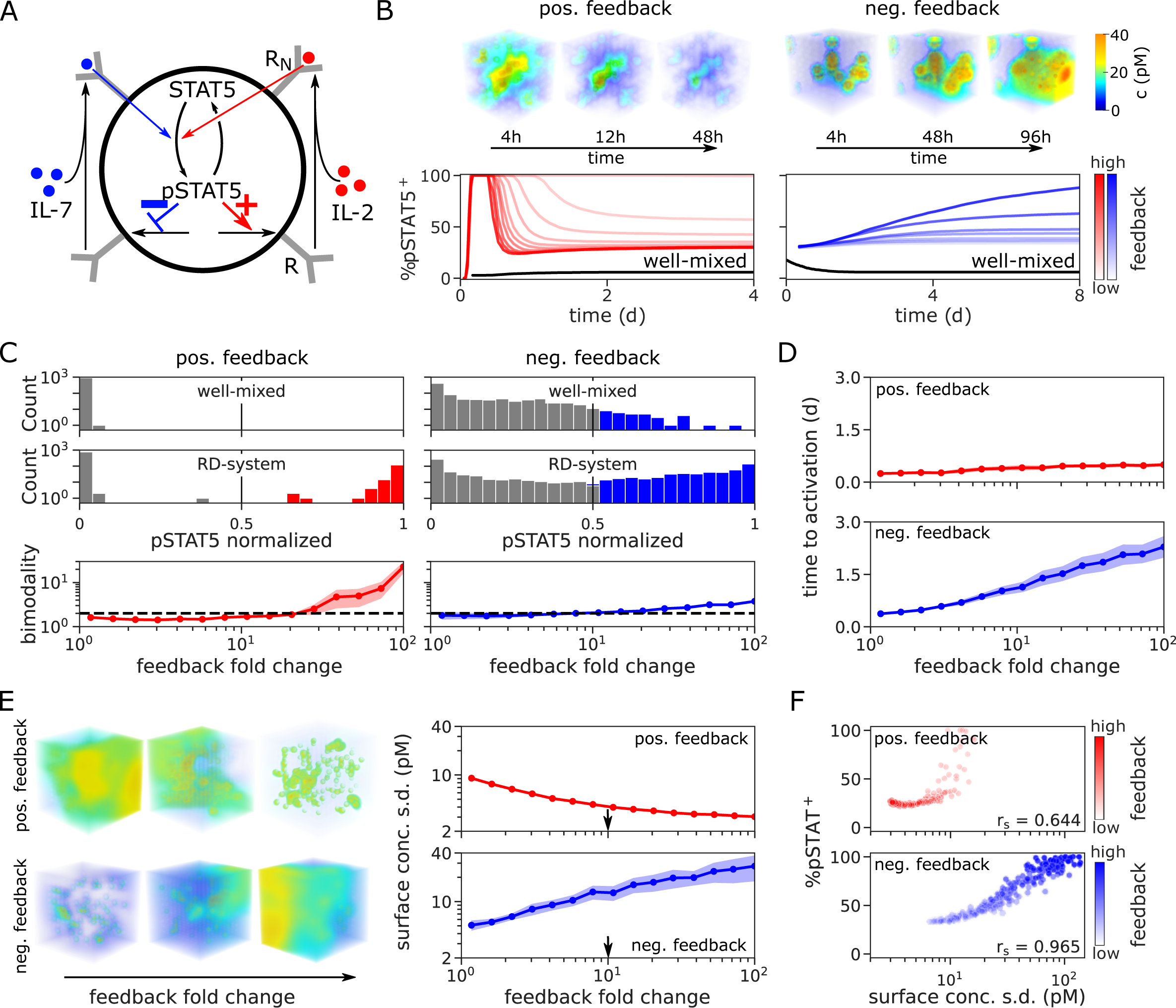
Feedback on receptor expression generates complex spatio-temporal dynamics. (**A**) Model scheme for positive and negative feedback on cytokine receptor expression. The pSTAT signal downstream of paracrine cytokine signaling induces either an increase (positive feedback, “IL-2”) or a reduction (negative feedback, “IL-7”) of receptor expression. (**B**) Kinetic simulation of the model illustrated in panel A, see text for details. Shown are the cytokine concentration field at three different time points (top), and the paracrine signaling efficacy measured by fraction of pSTAT5+ cells at varying feedback fold change ɣ (bottom). (**C**) Analysis of bimodality in cytokine receptor expression for positive and negative feedback. Shown are histograms of the pSTAT5 signal across cells (top), and the resulting bimodality as measured by Ashman’s D (bottom), at the last time point. Grey bars represent cells falling below a threshold value of 0.5 for pSTAT5, and dashed lines indicate a threshold value of 2 for Ashman’s D indicating significant bimodality. (**D**) Time until cells that are pSTAT5+ at steady-state (cf. panel C) reach the threshold value of 0.5 for the first time. (**E**) Analysis of spatial cytokine inhomogeneity with respect to receptor feedback. Shown are cytokine concentration fields (left) and corresponding values of the spatial s.d. in dependence of the feedback fold change ɣ (right), for positive and negative feedback as indicated. Vertical arrows indicate standard parameter values. (**F**) Correlation between the fraction of pSTAT^+^ cells and the spatial s.d. depending on feedback fold change. Each dot represents a single run. rs is the computed spearman’s rank correlation coefficient. Lines with shaded regions (panels C and E) represent average and SEM across 20 runs of a simulation.

In the IL-2 scenario, upon initializing cytokine secretion, the system quickly reached a transient state of high systemic cytokine concentration accompanied by increased STAT5 signaling activity and a subsequent increase in IL-2 receptor expression (Figure 3B and Figure S4B-D left panels), in agreement with experimental data^30^. To analyze the effects of negative feedback by means of the homeostatic IL-7 signal in comparison to positive feedback, we considered a thought experiment where the system is initially deprived of cytokine (Figure 3B and S4B-D, right panels). Upon initializing cytokine secretion, responding cells decreased receptor levels due to negative feedback, resulting in increased levels of cytokine concentration and STAT5 signaling. In the corresponding well-mixed simulations (Figure 3B and Figure S4C), cytokine concentrations remained below threshold for signal induction in both the IL-2 and the IL-7 scenarios.

Next, we sought to analyze the properties of positive and negative feedback in more detail. Generally, the population of responding cells can be categorized into two groups: (i) cells capable of maintaining a high level of STAT5 signaling activity, and (ii) cells which are unable to receive sufficient signal after an initial transient (Figure S4E-G). The emerging bimodal distribution for positive feedback is not present in well-mixed scenarios and shows a strong dependence on the feedback fold change (Figure 3C, left panel). On the other hand, negative feedback induced a more gradual response for both the well-mixed and RD-system, with the RD-system yielding a higher fraction of cells exhibiting stable STAT5 signaling activity (Figure 3C, right panel). Next to those differences regarding STAT5 distributions, also the time to activation shows marked differences between positive and negative feedback regulation (Figure 3D), with negative feedback showing a much slower response time that is subject to modulation by feedback fold change. Quite interestingly, strong positive feedback induced a decay and strong negative feedback induced an increase in measures of spatial cytokine inhomogeneity (Figure 3E and S4H-I), due to opposed effects on signal localization. This change in signal localization results in a similar change in cytokine signaling efficacy (Figure 3F), which is in line with the notion of the IL-7 receptor as an ‘altruistic’ signaling mediator^39^, as IL-7 uptake is stopped upon signal reception, so that cells further away from the cytokine-secreting cell are able to receive cytokine molecules.

In conclusion, we found the introduction of dynamic feedback to be a crucial control mechanism in shaping not only the distribution and timing of tissue responses but also the spatial cytokine gradients and signaling efficacy.

### Regulatory properties of the cytokine signaling niche

To understand the intertwined regulation of paracrine signaling via spatial inhomogeneities and feedback regulation in more detail, we shifted our focus to the immediate neighborhood of each secreting cell, which has previously been referred to as signaling niche^16^. Based on the results on feedback-driven spatial inhomogeneity (cf. Figure 3E), we hypothesized that positive feedback restricts the signaling range and negative feedback amplifies the signaling range, thus potentially enabling effective signaling towards a larger group of responder cells.

To identify the signaling niche of an individual cytokine secreting cell, we utilized density-based spatial clustering of applications with noise (DBSCAN) and defined a signaling niche as a cluster that contains at least one cytokine secreting cell and at least one activated cell (Figure 4A and B). Hence, the maximum possible number of niches equals the number of secreting cells (black line in Figure 4B), and since one cytokine-secreting cell is insufficient for effective paracrine signaling in the physiological parameter regime, the realized number of niches in the system typically falls far below that maximal number. Notably, we found that positive feedback leads to increased signaling activity inside a niche, and negative feedback to a higher fraction of activated cells outside the niche (Figure 4C), in line with our hypothesis on the signaling range.

**Figure 4:**
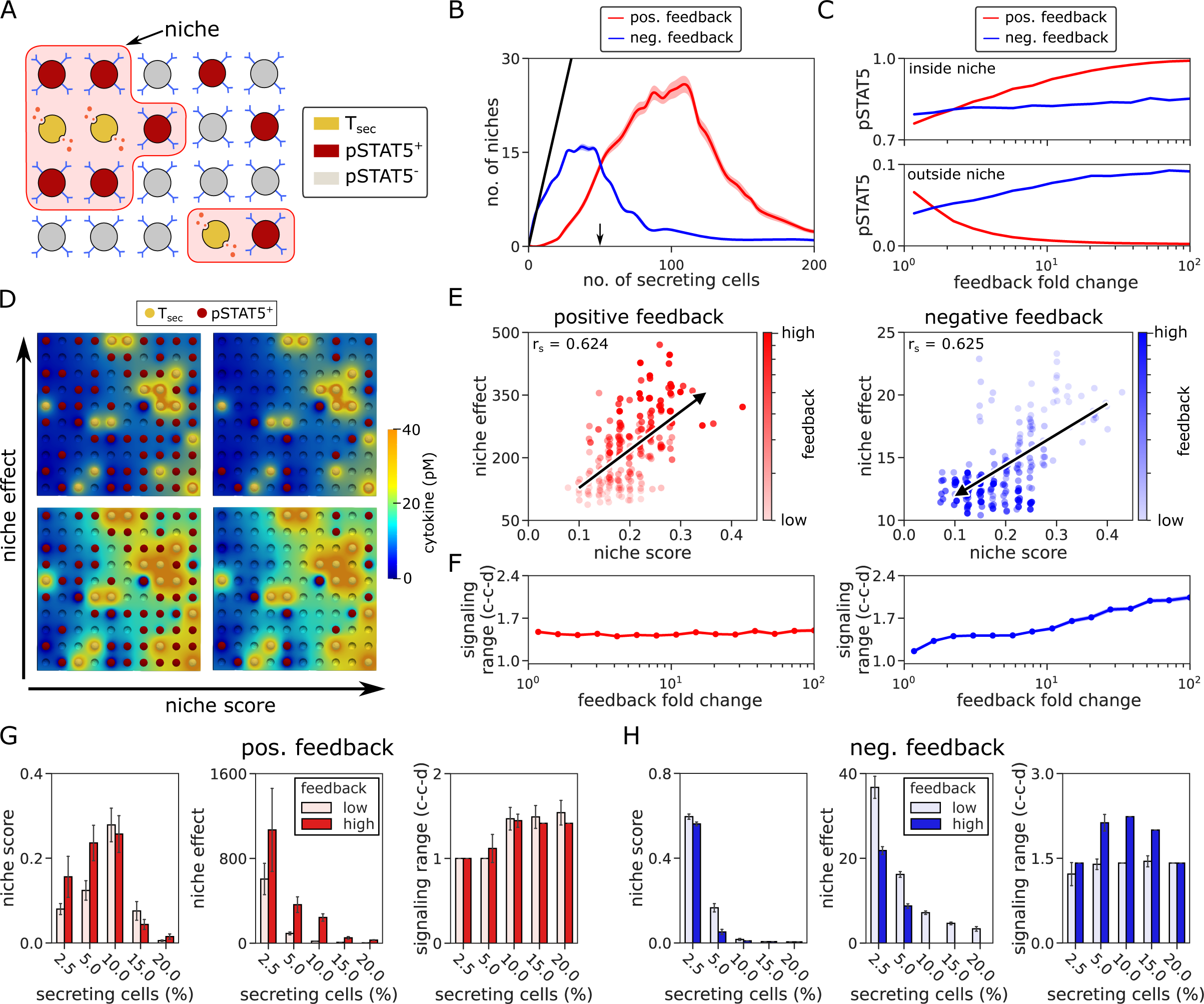
Niche score and niche effect characterize spatial patterning of paracrine cytokine signals. (**A**) Schematic illustrating the definition of a cytokine signaling niche: upon clustering of cells by theirpSTAT5 values, a niche is defined as a cluster that contains at least one secreting cell and one pSTAT5+ cell. (**B**) Number of niches (cf. panel A) in dependence of the number of cytokine secreting cells. Black line: theoretical maximum (every Tsec cell forms a separate niche). Black arrow: standard parameter (cf. Table 1). (**C**) Average pSTAT5 values of cells inside and outside of a cytokine signaling niche. (**D**) Visualization of the quantities niche score and niche effect. For visualization purposes 15% secreting cells were used. The niche score is defined as the number of niches divided by the number of secreting cells in the system, the niche effect as the fraction of pSTAT5+ cells within the niche divided by outside of the niche. See also Figure S5A. (**E**) Correlation between niche score and niche effect for positive and negative feedbackThe black line shows a linear fit of all data points. The arrow head was added to indicate the direction of increased feedback. rs is the computed spearman’s rank correlation coefficient. (**F**) Signaling range in dependence of feedback fold change. (**G-H**) Niche score, effect and signaling range for varying fractions of cytokine secreting cells. The bar color indicates feedback fold-change, low as ɣ = 2 and high as ɣ = 100, cf. panels C andF. Lines with shaded regions (panels B, C, F) and errorbars (F and G) indicate average and SEM across 20 runs of the simulation.

To proceed to a more direct quantitative analysis, we defined the ‘niche score’ as the ratio between the number of niches and the number of secreting cells, and the ‘niche effect’ as the ratio between the average pSTAT5 signal inside and outside of the niche (Figure 4D-E and S5A). A high niche score indicates well-separated niches, and a high niche effect indicates that the signal is primarily located within the niche compared to outside, in other words, it accounts for the leakiness of the cytokine niche. Furthermore, our definition of the signaling niche allowed to quantify the signaling range as average distance between cytokine-secreting cells and maximal-distance responder cells within a niche (Figure 4E). The niche score and niche effect in conjunction with the signaling range (Figure 4F) revealed that positive feedback enhances niche separation (high niche score) and induces a highly localized signal within each niche (high niche effect with low signaling range), while negative feedback causes niches to merge (low niche score) and increases cytokine leakiness (low niche effect and high signaling range).

Those system properties further depend on the number of secreting cells, with small numbers limiting the effect of both positive and negative feedback, since only few responder cells are activated (Figure 4G and S5B), and high numbers additionally dampening the effect of negative feedback (Figure 4H and S5C). That latter effect can be attributed to larger niches requiring longer signaling ranges, reflecting the previously detected (cf. Figure 3D) increase in activation time under negative feedback.

Overall, our detailed quantitative analysis of the signaling niche revealed that local receptor expression feedback is able to influence not only signaling efficacy but also signaling range and niche separation, facilitating an adaptive response to variable stimulus intensities.

### The local tissue architecture provides an additional regulatory layer for spatiotemporal cytokine signals

Thus far, to systematically investigate the spatiotemporal dynamics, cells were placed randomly on a cubic grid in all simulations shown. To analyze the effect of additional constraints imposed by the local tissue architecture, we also designed a grid-free simulation setup allowing to induce clustering of specific cell types in a tightly controlled manner, by variation of the clustering strength φ. That clustering strength correlates well with the silhouette and Calinski-Harabasz scores, which are established metrics for clustering quality (Figure S5D). Such clustering of specific cell types is a wide-spread property of immune-cell populations and can be mediated by chemokine signals or also physical barriers, such as imposed by stromal cells in the lymph node^40,41^.

Here, we initially considered a situation where IL-2 secreting T cells accumulate in the vicinity of a specific antigen-presenting cells presenting their cognate antigen (Figure 5A). As expected, such co-localization of cytokine-secreting cells imposes a substantial increase in paracrine signaling efficacy in our simulations, since locally enriched cytokine concentrations levels allow to overcome the activation threshold in nearby cells. For low-to-moderate numbers of cytokine-secreting cells, quantitative analysis confirmed that effect in terms of the number of activated cells, the niche score defined above (Figure 5B) and the signaling range, despite an overall reduction of IL-2 concentration levels (Figure S5E). Quite interestingly, at very high numbers of cytokine-secreting cells, co-localization can also have the opposite effect and induces a reduction in the number of activated cells (Figure 5B, inset), which can be explained by the isolating effect of nearby responding cells preventing long-range paracrine signaling. To provide immune tolerance and reduce the risk of immune-responses to self-antigen, paracrine IL-2 signaling is mitigated by regulatory T (Treg) cells that express large numbers of high-affinity IL-2 receptor and can act as strong cytokine sinks^15,19,42^. In particular, it has been suggested that Treg cells can effectively take up IL-2, because they are stimulated by cognate antigen presented by the same antigen-presenting cell as the corresponding effector T cells and thus are co-localized in the same spatial signaling niche^18,43^ (Figure 5C, left panel). Indeed, we found that the inhibitory effect of Treg cells is rather limited if they are placed randomly, that is φ=0 (Figure 5C-D). At least a partial co-localization of Treg cells with cytokine-secreting cells seems required for effective reduction of cytokine signal and signaling range (Figure S5F), and can induce almost complete signal inhibition already at a fraction of only 4% Treg cells in our simulations.

**Figure 5:**
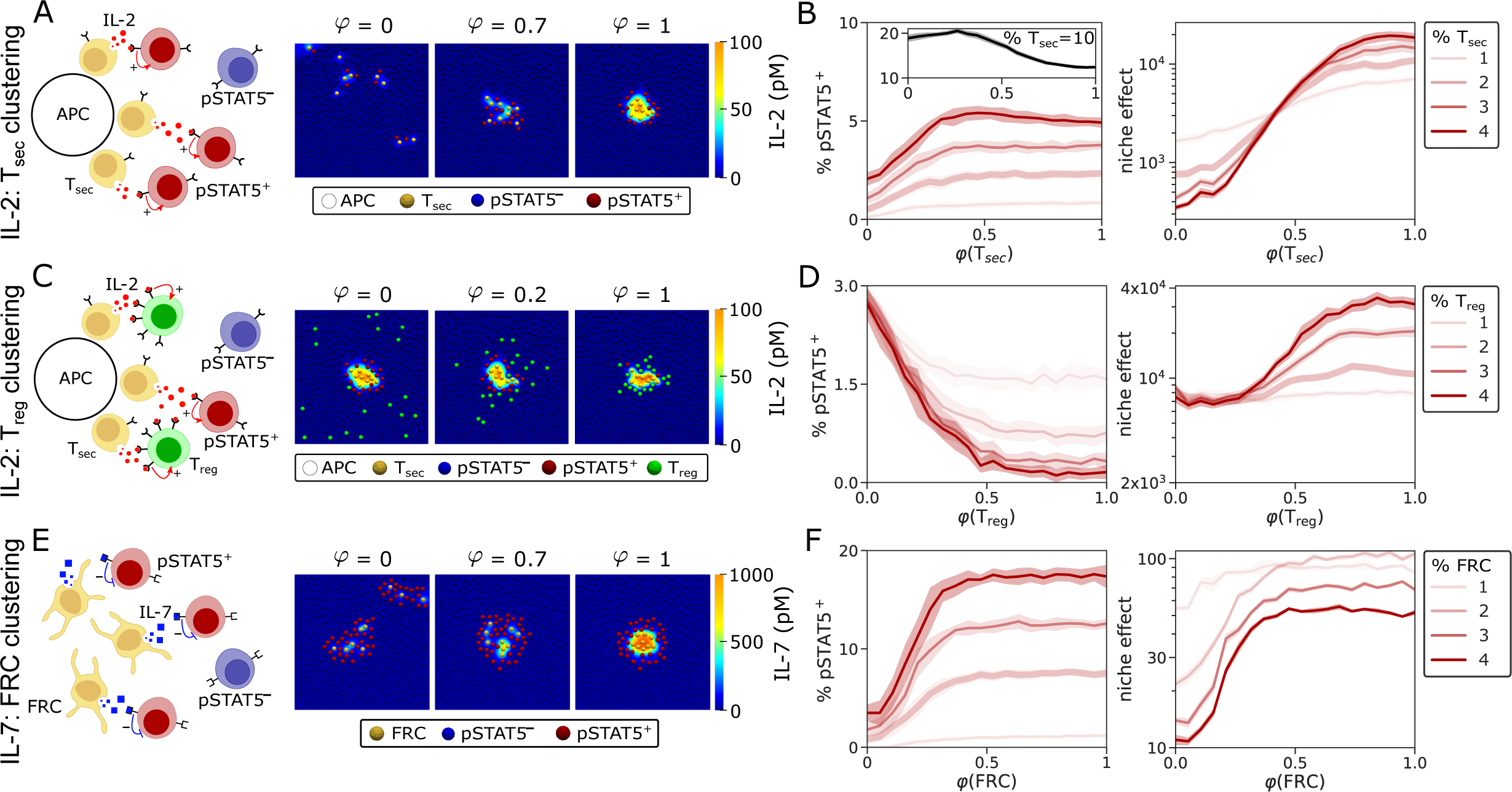
Spatio-temporal cytokine dynamics in the context of an established tissue architecture. (**A**) Schematic (left panel) and typical simulations (right panel) of a scenario with IL-2 secreting T cells (Tsec) in the vicinity of an antigen presenting cell (APC), surrounded by responder T cells. φ: clustering strength of Tsec cells. Shown are two-dimensional snapshots for the purpose of visualization, the analysis below is performed on full 3D-simulations. (**B**) Analysis of the fraction of pSTAT5+ cells (left) and the niche effect (right) in dependence of the clustering strength φ for the scenario visualized in panel (A). Lines with shaded regions indicate average and SEM across 20 runs of each simulation. (**C-D**) Regulatory T cells (Treg) expressing high numbers of IL-2R are co-localized with Tsec cells in the vicinity of an antigen-presenting cell. See panels (A-B) above for details. (**E-F**) Fibroblastic reticular cells (FRC) secrete IL-7, which is taken up by naive T helper cells.

While IL-2 signaling in the local environment created by antigen-presenting cells facilitates fast and unambiguous decision-making on T cell activation, it has been argued that IL-7, a cytokine essential for T cell survival, should be distributed more widely within the system^39^. Our results on the spatio-temporal effects of negative feedback (cf. Figure 4E) supported that claim, as we observed an increase in paracrine signaling efficacy and spatial range. To investigate that phenomenon in a more specific scenario, we introduced co-localization of IL-7 secreting cells analogous to the IL-2 case (Figure 5E). As anticipated, we found that increased clustering of cytokine-secreting cells not only improved signaling efficacy (Figure 5E-F), but also caused an increase in cytokine concentration levels and signaling range (Figure S5G).

Hence, we found that the spatial composition of cell types with specific properties can provide another layer of control over the range and efficacy of paracrine cytokine signaling, acting together with the amount of cytokine secreting cells, the distribution of cytokine receptor expression and cytokine-induced feedback regulation.

## Discussion

Cell-cell communication using diffusible ligands is a widespread mechanism to exchange information in multi-cellular organisms, and is particularly important in the collective decision-making processes of the mammalian immune system. Compared to intracellular processes, the larger spatial domain of such cell-cell interaction dynamics increases the potential for inhomogeneous distributions and non-intuitive spatial patterning^44,45^ of signaling mediators. Nevertheless, the high diffusivity (∼10 µm^2^/s) of small proteins such as cytokines suggests that concentration inhomogeneities may disappear very rapidly on the relevant time- and length scales, that is cell-cell distances in lymph nodes (<5 µm) and times for signal integration and cell-differentiation (minutes to hours). By systematic analysis of physiological scenarios of cytokine signaling using an efficient finite-element simulation setup, we found that spatial cytokine inhomogeneities do not arise by diffusion processes per se. Rather, additional factors are required, such as a sparse occurrence and all-or-none behaviour of cytokine secreting cells or heavily skewed distributions of cytokine receptors across cells. Thus, concentration inhomogeneities are essentially a property that is under control of the cell population.

Our mathematical formulation combines a description of signal processing on the cellular level with a biophysical description of cytokine diffusion. The signaling range of cytokine-secreting cells is diffusion-limited, since cytokine uptake requires diffusion of the cytokine through extracellular space, in addition to binding to its high-affinity receptor^46,47^. The importance of this diffusion limit becomes apparent when considering the diminished signaling efficacy in our well-mixed model implementation. In the reaction-diffusion model, considering non-linear uptake dynamics on the surface of responding cells, we found that both the fraction of secreting cells and heterogeneity in receptor expression were able to generate cytokine concentration inhomogeneities. Previous studies have indicated values between 1% and 20% for the fraction of cytokine-secreting cells^14,25^ and a coefficient of variation for the receptor distribution that is close to 1^4,24^. Under those conditions, our results suggest a higher contribution of the fraction of cytokine-secreting secreting cells to spatial inhomogeneities within the physiological parameter range.

Another important factor shaping cytokine signaling dynamics is feedback regulation of receptor expression on responding cells, which has been reported not only for IL-2 and IL-7 as the key interest of this study, but also for other cytokines including IL-4, IL-21, TGF-β and IFN-γ^5,31–33,48–50^. It is known that positive feedback can induce an all-or-none type of response to a signal, while negative feedback leads to more gradual and homogeneous dynamics and can have an effect on the time-scale of the response to an input signal^51,52^. In line with that and with previous studies^12^, we found that spatial cytokine gradients can induce significant bifurcations in activation patterns of the naive T cell in the presence of positive feedback mediated by the cytokine IL-2. Interestingly, that model behavior does not occur in a parallel well-mixed scenario, highlighting the requirement of spatial cytokine distributions for paracrine stimulation even in the presence of positive feedback. Considering IL-7 as a cytokine promoting negative feedback on receptor expression, we observed an increase in signaling range that propagates with time, resembling a signaling wave that spreads through the cell population. That phenomenon is well in-line with the notion of IL-7 responder cells as ‘altruistic’ ^39^, since they provide access to IL-7 to nearby cells by downregulating their receptors.

To quantify the localization and effectiveness of cytokine signaling within the microenvironment around cytokine-secreting cells, we propose the niche score and niche effect as summary statistics that may also serve for comparison with multi-color histology data in future research. The niche score offers an understanding of how well separated niches are, while the niche effect quantifies how effectively cytokine signal is localized within each niche. Utilizing these spatial statistics, we found that the up-regulation of receptors in the IL-2 scenario results in a localization of activation and signal within a niche. We identified the opposite behavior in the case of IL-7 signaling, where downregulation of receptors delocalizes the signal, thus inducing a more homogeneous cytokine concentration field. Consequently, paracrine signaling efficacy depends more on the total amount of cytokine molecules in the system rather than its location, which is in line with the biological function of IL-7 as a survival signal controlling the size of a cell population in homeostasis.

Finally, we asked how the local tissue architecture in terms of already established clustering and co-localization of specific cell types would modulate signaling efficacy and spatial cytokine patterning. Our model simulations revealed that the spatial clustering of cytokine-secreting cells can substantially increase signaling efficacy, as cytokine secretion by multiple cells in a signaling niche combines to act on responder cells. Moreover, we found that co-localization of cytokine-secreting cells and cells with a high capacity of cytokine uptake, such as described for Treg cells in the context of IL-2 signaling^18^, is a highly efficient mechanism to control the effectivity of paracrine signals and thus modulate the degree of immune tolerance.

Overall, we found that the spatial relationships and individual properties of cytokine-secreting cells and cells expressing high-affinity cytokine receptor species can critically regulate the efficacy of cell-communication. Future research may combine our approach with quantitative models on germinal-center dynamics^53–55^ and multiplexed histology data^56–58^ characterizing tissue organization in lymphoid organs and the tumor micro-environment, paving the way to a unified, quantitative understanding of spatio-temporal decision-making in immune-cell populations.

## Acknowledgments

This work was supported by the German Research Association (TH 1861/5-1), Germany’s Excellence Strategy (EXC2151-390685813 and EXC2047-390873048), and the Leibniz Association (Best-minds junior research group), all to K.T.

## Methods

### Software, simulations and statistics

All simulations are carried out using the finite-element solver FEniCS^59^ with P1 elements, a generalized minimal residual methods (GMRES) solver and algebraic multi grid (AMG) preconditioner. The solution accuracy is controlled by the Krylov solver tolerance for linear and the Newton solver tolerance for the non-linear boundary conditions. For a description of the weak formulation of the models used therein, refer to the Supplementary Text. Standard parameter values are listed in Table 1.The extracellular space was discretized as a uniform tetrahedral mesh using Gmsh^60^, the mesh fidelity was chosen to yield a mesh with approximately 100,000 degrees of freedom in a cube of 240 µm edge length. To minimize boundary effects the outermost layer of cells is disregarded in our analysis.

The ODE model was solved using a SciPy ODE solver. All parameter values are listed in Table 1. The standard deviation (SD) was computed using the surface concentration of all cells in one configuration. The gradient was computed through the average norm of the cytokine field gradient. The Spearman rank-order correlation coefficient (rs) was computed using SciPy. To quantify the bimodality of a distribution we calculated the relative separation using Ashman’s D^61^. The signaling range was determined using the distances of responding cells to their closest secreting cell inside all niches and calculating the 0.05^th^ largest percentile.

## Mathematical Models

### Core-model of spatially resolved cytokine dynamics

Let Ω be the extracellular space, Θ the outer boundary of that space and Γ_i_, *i* = 1, …, *N*_cells_, denote the cell-surfaces. In all simulations, for each cytokine considered, we assumed the cytokine concentration *c*(*x*), *x* ∈ Ω to be in quasi-steady-state, and we imposed no-flux conditions at the outer boundary, that is 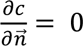. on Θ Further, assuming homogeneous cytokine secretion and uptake on each cell-surface with area *A*_*i*_, the interaction between a cell and the extracellular cytokine concentration is realized through generalized Robin boundary conditions (cf. Ref.^12,13^):

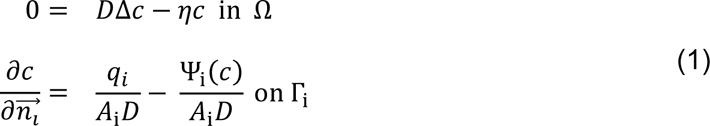

Here, *D* is the diffusion coefficient, η is a homogeneous decay rate and Δ is the three-dimensional Laplace operator. For the sake of simplicity, we assume a uniform cell-size *A*_i_: = *A*_cell_ throughout. Furthermore, we take the secretion rate *q*_*i*_ and the surface concentration as average values across the cell-surface for each cell, that is 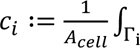 *c ds*. Hence, the uptake rate takes the form

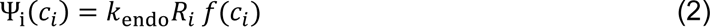

where *R*_*i*_ denotes the number of receptors expressed on the cell-surface. In the case of linear cytokine uptake, the uptake function takes the form 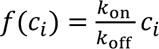, where *k*_on_ is the cytokine-receptor association rate^12,13^. A more realistic description of cytokine uptake takes into account that the uptake capacity of a cell is limited not only by the amount of receptors expressed on the surface, but also by the intracellular machinery for cytokine degradation and ‘recycling’ of cytokine receptors^12,62,63^. That results in an uptake function of the form 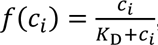, where *K*_D_ is the half-saturation constant for cytokine uptake. We found that this form of the uptake function can be justified by a mechanistic, Michaelis-Menten type description of cytokine-binding to its receptor (Supplementary Text, “Description of uptake dynamics”). Further, assuming *k*_endo_ = *k*_off_, we retrieve the canonical form of the linear uptake function Ψ_i_(*c*_*i*_) = *k*_on_*R*_*i*_ *c*_*i*_. In the case of non-secretory responder cells, we set *q*_*i*_ = 0 and *R*_*i*_ = *R_resp_* in Equation 1, whereas we take *q*_*i*_ = *q*_*sec*_ and *R*_*i*_ = *R*_*sec*_ for cytokine-secreting cells. The latter ones express lower numbers of cytokine receptors and thus exhibit a reduced uptake rate Ψ_i_(*c*_*i*_) (cf. Table 1). Moreover, we account for receptor heterogeneity on responding cells (Figure 2) by sampling *R*_*i*_ from a log-normal distribution with mean *R*_*resp*_.

As a primary output of our model simulations, we assess cellular activation in terms of the fraction of phosphorylated STAT5, which we take as

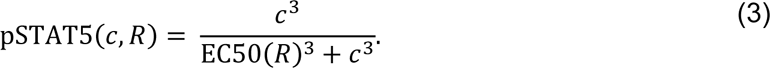

A cell is considered activated if its pSTAT5 level reaches values pSTAT5(*c*_*i*_, *R*_*i*_) ≥ 0.5. In line with experimental data^24^, we account for a dependency of the EC50 value on the level of receptor expression through 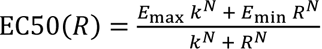, where *E*_max_ *E*_min_, *k* and *N* are parameters determined through fitting of experimental data^24^. Model parameters (Table 1) could in many cases be assigned to or derived from published experiments.

### Receptor feedback kinetics

In order to analyze the delayed receptor feedback introduced in Figure 3, we considered a time-dependent change in receptors, leading to a time-dependent cytokine distribution. That is, our quasi-stationary diffusion problem, Equation 1, is generalized to a series of such problems, via Ψ_i_(*c*_*i*_) = Ψ_i_(*c*_*i*_(*t*), *R*_*i*_(*t*)) in Equation 2, given by a system of ODE for receptor expression in each cell:

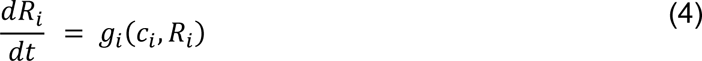

Specifically, we chose a receptor feedback function

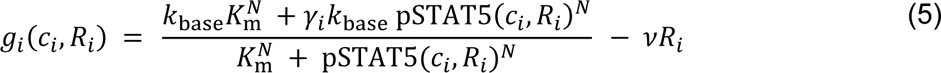

that depends on the cellular pSTAT5 level, the minimal receptor production rate *k*_base_, the half saturation value for activation *K*_m_ and the receptor decay rate *v*. The feedback fold change γ_*i*_ depends on the cell type, here we take γ_*i*_ = 1 for cytokine-secreting cells, γ_*i*_ ɛ (1, 100] for positive and γ_*i*_ ɛ [0.01, 1) for negative feedback on responding cells. For visualization, to allow for a unified x-axis, the fold change for negative feedback was inverted. To account for delayed regulation of receptor expression on the responding cell *i* caused by intracellular processes such as signal transduction and gene expression, we supplemented Equation 4 by equations for auxiliary states (‘linear chain trick’, cf. Figure S4A), as previously described^37^:

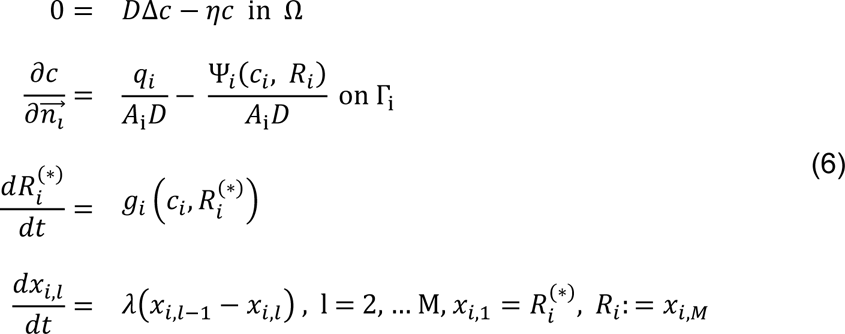

### Well-mixed model

In the well-mixed case, i.e. assuming infinitely fast diffusion, the homogeneous extracellular cytokine distribution is described by an ODE. Namely, by neglecting the cell volumes, the core PDE model (Equation 1) converges for *D* → ∞ to an ODE in terms of the system-wide quantities *q*_*tot*_: = *q*_*sec*_*N*_*sec*_, where *N*_*sec*_ denotes the number of secreting cells, and ψ_*tot*_(*c*) ≔ *k*_endo_*R*_*tot*_*f*(*c*) with *R*_*tot*_ ≔ *N*_*sec*_*R*_*sec*_ + *N*_*resp*_*R*_*resp*_, considered at steady-state (Supplementary Text, “Deriving the well-mixed model”)^64^:

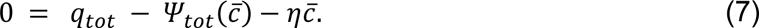

As in the core PDE model, we have 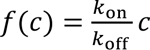 for linear uptake and 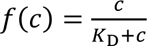 for saturated uptake. Parameters were chosen identically to the core model and are listed in Table 1.

Furthermore, the steady-state solution 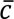 of the well-mixed ODE is the average of the cytokine distribution *c* described by the core PDE^64^, i.e. 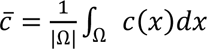. That allows for a direct comparison to the core model in terms of analytical expression of values for the average cytokine concentration (Supplementary Text).

Analogous to the core PDE model, we introduced delayed receptor feedback in the well-mixed model by a time-dependent uptake function ψ_*tot*_(*c*) = ψ_*tot*_(*c*(*t*), *R*_*tot*_(*t*)) and receptor expression dynamics 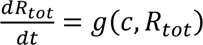.

### Random cell positioning

To achieve an efficient 3-dimensional random cell positioning (cf. Figure 5), we utilize Bridson sampling, a Poisson disk sampling approach^65^. The average cell-cell distance and the amount of sampling steps are chosen to yield the same average cell density as in the grid approach employed in all other simulations (Figure 1-4).

### Quantifying spatial patterns

The surface concentration average refers to the average concentration over the surface of all responding cells in the system, the surface concentration s.d. refers to the standard deviation of the concentration of all responding cells. The gradient refers to the sum of the cytokine gradient in the extracellular space. To calculate the niche score and niche effect in Figure 4 it is necessary to calculate which pSTAT5^+^ cells are being activated by which secreting cell. This is achieved by running the DBSCAN algorithm with all pSTAT5^+^ and secreting cells spatial coordinates as input and the average cell-cell distance, typically 20 µm, as the maximum distance between two samples. This algorithm assigns each cell to a cluster, which is a group of cells, based on the spatial density of cells. Any resulting clusters not containing at least one pSTAT5^+^ and one secreting cell were disregarded, which only happens in edge cases since activation is localized around secreting cells.

### Clustering formalism

In order to achieve the clustering of secreting cells around APC in Figure 5 we developed an algorithm that generates clusters of cell types around a set of user-defined points in space called clustering points. Each cell type is represented by a fraction and a clustering strength, which determines how clustered the cells of each type will be around the clustering points. The algorithm uses a kernel density estimation (KDE) to assess the probability of each possible cell position being part of a cluster. It then samples cell positions based on these probabilities to assign each position a cell type.

## Notes

### Competing Interest Statement

The authors have declared no competing interest.

### Summary of Updates

Small correction in Figure 1.

