## Supplemental Text and Figures for "Diffusion-limited cytokine signaling in T cell populations"

### 1 Supplementary Text

#### Deriving the weak form

In order to numerically solve the partial differential equation (PDE) posed in Equation 1 in the main text, one first has to derive the weak form. Let $\Omega \subset \mathbb{R}^3$ be the extracellular space, $\Theta \subset \mathbb{R}^2$ be the outer boundary of that space and $\Gamma_i \subset \Omega, \Gamma_i \subset \mathbb{R}^2$ the closed boundaries for every cell.

$$0 = D\Delta c - \eta c = D\nabla^2 c - \eta c \tag{1}$$

$$\frac{\partial c}{\partial \vec{n}} = 0 \text{ on } \Theta \tag{2}$$

$$\frac{\partial c}{\partial \vec{n}_i} = \frac{q_i}{A_i D} - \frac{\Psi_i(c_i)}{A_i D} \text{ on } \Gamma_i. \tag{3}$$

To derive the weak formulation necessary to solve this problem, we multiply Equation (1) by a test function $v$ and integrate over the extracellular space $\Omega$, yielding

$$D \int_\Omega (\nabla^2 c) v \, dx = \int_\Omega \eta c \cdot v \, dx \tag{4}$$

with $dx$ as the differential element for the integration over $\Omega$. Utilizing integration by parts and the divergence theorem, we can write the left side of this equation as

$$D \int_\Omega (\nabla^2 c) v \, dx = -D \int_\Omega \nabla c \cdot \nabla v \, dx + D \int_{\partial\Omega} \frac{\partial c}{\partial \vec{n}} v \, ds \tag{5}$$

with $ds$ as the differential element for the integration over all boundaries of $\Omega$. Using Equation (2) and (3) the integration over the boundary $\partial\Omega$ can be written as

$$\begin{aligned}\int_{\partial\Omega} \frac{\partial c}{\partial \vec{n}} v \, ds &= \int_\Theta \underbrace{\frac{\partial c}{\partial \vec{n}}}_{=0} v \, ds + \sum_i \int_{\Gamma_i} \frac{\partial c}{\partial \vec{n}_i} v \, ds = \sum_i \int_{\Gamma_i} \frac{1}{A_i D}(q_i - \Psi_i(c_i)) v \, ds \\ &= \sum_i \frac{1}{D}(q_i - \Psi_i(c_i)) v.\end{aligned} \tag{6}$$

With this we get the weak form

$$D \int_\Omega \nabla c \cdot \nabla v \, dx = -\int_\Omega \eta c \cdot v \, dx + \sum_i (q_i - \Psi_i(c_i)) v \tag{7}$$

which we use in FEniCS to solve the RD-system.

#### Deriving the well-mixed model

To evaluate the spatial effects in the PDE based core model we compare our results with the ordinary differential equation (ODE) based well-mixed model (cf. Figure 1 in the main text). The ODE approach does not include spatial information but rather assumes that the extracellular volume is well-mixed. Mathematically this can be achieved by neglecting the cell volumes, leading to cytokine secretion and uptake to be considered system-wide with respective rates

$$q_{\text{tot}} = D \sum_{\text{sec. cells } i} \int_{\Gamma_i} \frac{q_i}{A_i D} \, ds = q_{\text{sec}} N_{\text{sec}} \tag{8}$$

$$\Psi_{\text{tot}}(c) = D \sum_{\text{cells } i} \int_{\Gamma_i} \frac{\Psi_i(c)}{A_i D} \, ds = \sum_{\text{cells } i} \Psi_i(c) = \begin{cases} k_{\text{on}} R_{\text{tot}} c & \text{linear} \\ k_{\text{endo}} R_{\text{tot}} \frac{c}{K_{\text{D}} + c} & \text{saturated.} \end{cases} \tag{9}$$

Thus, the core PDE in Equation (1) with neglected cell volumes can be written as

$$0 = D\Delta c - \eta c + q_{\text{tot}} - \Psi_{\text{tot}}(c) \text{ in } \Omega \tag{10}$$

$$\frac{\partial c}{\partial \vec{n}} = 0 \text{ on } \Theta. \tag{11}$$

Using Gronwall estimations, it was shown that for $D \to \infty$, the solution $c$ of Equation (10) converges exponentially to its space average $\bar{c} = \frac{1}{|\Omega|}\int_\Omega c(x)\ dx$ [S1, Thm 3.1]. This yields the well-mixed ODE at steady-state, solving Equation (8) in the main text as

$$\bar{c} = \frac{q_{\text{sec}} N_{\text{sec}}}{k_{\text{on}} R_{\text{tot}} - \eta} \tag{12}$$

in case of the linear uptake and

$$\bar{c} = \frac{-q_{\text{tot}} + k_{\text{endo}} R_{\text{tot}} + K_{\text{D}}\eta - \sqrt{K_{\text{D}}^2\eta^2 + 2K_{\text{D}}\eta(q_{\text{tot}} + k_{\text{endo}} R_{\text{tot}}) + (q_{\text{tot}} - k_{\text{endo}} R_{\text{tot}})^2}}{2\eta} \tag{13}$$

for the saturated uptake.

#### Description of uptake dynamics

##### *Saturated Uptake*

To account for saturating effects in our uptake dynamics, we employ classical Michaelis-Menten kinetics in order to find the uptake function $\Psi$ described in the methods in the main text. The binding of cytokine $c$ to a receptor $R$, formation of a cytokine/receptor complex followed by internalization can be written as

$$c + R \underset{k_{\text{off}}}{\overset{k_{\text{on}}}{\rightleftharpoons}} C \xrightarrow{k_{\text{endo}}} P \tag{14}$$

with $C$ as cytokine/receptor complexes and P as the internalized cytokine/receptor complexes with their corresponding rates $k_{on}$ for binding, $k_{off}$ for unbinding, and $k_{endo}$ for endocytosis.

From this, we can define the differential equations as

$$\frac{dR}{dt} = k_{\text{off}} C - k_{\text{on}} R c + a \tag{15}$$

$$\frac{dC}{dt} = k_{\text{on}} R c - k_{\text{off}} C - k_{\text{endo}} C \tag{16}$$

with $a$ as a base receptor production.

Under the assumptions that $c >> R$ and $P_0 = 0$, the uptake flux $\Psi$ can be considered as the velocity at which C is converted into P, resulting in

$$\Psi = k_{\text{endo}} C \tag{17}$$

Assuming rapid equilibrium, which means

$$k_{\text{on}} R c = k_{\text{off}} C \tag{18}$$

and employing mass conservation $R = R_0 - C$, where $R$ is the number of unbound receptors on the cell and $R_0$ is the overall amount of receptors, we get

$$k_{\text{on}}(R_0 - C)c = k_{\text{off}} C \tag{19}$$

$$k_{\text{on}} R_0 c = (k_{\text{off}} + k_{\text{on}} c) C \tag{20}$$

$$\implies C = \frac{k_{\text{on}} R_0 c}{k_{\text{off}} + k_{\text{on}} c} = \frac{R_0 c}{\frac{k_{\text{off}}}{k_{\text{on}}} + c} = \frac{R_0 c}{K_{\text{D}} + c} \tag{21}$$

with this we can write Equation (17) as

$$\Psi(c, R_0) = k_{\text{endo}} C = k_{\text{endo}} R_0 \frac{c}{K_{\text{D}} + c}. \tag{22}$$

Here $\Psi(c, R)$ is linear in R and saturates via a Hill-function in $c$.

***Linear uptake***

To account for any effects introduced by the previously described saturated uptake dynamic, we derived an uptake function linear in $c$, which we use in Figure S3 to delineate saturating effects. Assuming $c << K_D$ with $K_\mathrm{D} = \frac{k_\mathrm{off}}{k_\mathrm{on}}$, we can then simplify (22) as

$$\Psi(c, R_0) = k_\mathrm{endo} R_0 \frac{c}{K_\mathrm{D} + c} = k_\mathrm{endo} R_0 \frac{c}{\frac{k_\mathrm{off}}{k_\mathrm{on}} + c} \approx k_\mathrm{endo} R_0 \frac{c}{\frac{k_\mathrm{off}}{k_\mathrm{on}}} \tag{23}$$

$$= \frac{k_\mathrm{endo}}{k_\mathrm{off}} k_\mathrm{on} R_0 c \tag{24}$$

If we further assume $k_\mathrm{endo} = k_\mathrm{off}$, it simplifies to

$$\Psi(c, R_0) = k_\mathrm{on} R_0 c \tag{25}$$

resulting in a function that scales linear with $c$.

***Transition from saturated to linear uptake***

As observed in Figure S3D, the saturated uptake dynamics can typically transition into a linear form if the condition $c << K_\mathrm{D}$ is fulfilled. However, this cannot be fulfilled by increasing $K_\mathrm{D}$ since $c$ depends on $K_\mathrm{D}$. To highlight this dependence, we can consider the saturated well-mixed case (cf. Equation 7 in the main text) with $\eta = 0$:

$$0 = q_\mathrm{tot} - k_\mathrm{endo} R_\mathrm{tot} \frac{c}{K_\mathrm{D} + c} \tag{26}$$

$$\implies c = \frac{q_\mathrm{tot} \cdot K_\mathrm{D}}{k_\mathrm{endo} R_\mathrm{tot} - q_\mathrm{tot}}. \tag{27}$$

Here $c$ is directly proportional to $K_\mathrm{D}$, so any increase in $K_\mathrm{D}$ will yield the same increase in $c$. To fulfill the initial condition, we decrease $q$, and hence $q_\mathrm{tot}$, yielding a linear decrease in $c$ since $k_\mathrm{endo} R_\mathrm{tot}$ remains constant. In the well-mixed model, our saturated uptake dynamics transition into the linear regime only during low cytokine secretion. In the spatial RD-system, the cytokine concentration $c$ depletes with increased distance to a secreting cell, resulting in the gradual transition from saturated to linear uptake.

#### Analytical approximation of the cytokine concentration field

The analytical approximation, taken from [S2] and used to verify our results in Figure S1 and S3, calculates the cytokine concentration for an environment with a high density of responding cells around a single secreting cell. It also assumes radial symmetry and is defined as

$$c(r) = \frac{q\rho}{k_\mathrm{on} r} \frac{4\pi D r (L + \rho) + k_\mathrm{on}(L - r + \rho) N R}{k_\mathrm{on} L R_\mathrm{sec} N R + 4\pi D \rho (L + \rho)(R_\mathrm{sec} + N R)} \tag{28}$$

with $\rho$ as the cell radius, $L$ as the synaptic distance, $N$ as the number of IL-2 consuming responder cells and $R_\mathrm{sec}$ as the receptors on secreting cells. Since the standard setup in our core model uses multiple cytokine secreting cells which are not necessarily isolated, this solution can only be used to approximate the cytokine concentration in this scenario.

#### 2 Supplementary Figure Legends

##### Figure S1, related to Figure 1 in the main text.

**(A)** Schematic of the code structure. Preprocessing, simulation and postprocessing are run independently. **(B)** Computation time for one run over increasing mesh fidelity, measured in mesh vertices (left) and over system size for a fixed mesh fidelity (right). **(C)** Average surface concentration over the amount of offset layers. An offset layer refers to a shell of cells, e.g. the first offset layer contains the outermost cells of the system. The vertical arrow indicates the standard parameter value. **(D-E)** Analysis of the RD- and well-mixed system against the analytical approximation in equation (28). Shown are (D) a setup with one secreting cell in the center where the analytical approximation holds and (E) a setup with 10% randomly distributed secreting cells where the analytical approximation does not hold. All other parameters are kept at standard values.

##### Figure S2, related to Figure 1 in the main text.

**(A-B)** Shown are the (A) average gradient over the extracellular space and (B) surface concentration s.d. over all responding cells as measures for inhomogeneity over the fraction of secreting cells (Tsec), simulations are designed in such a way that the total number of secreted cytokine molecules and the total number of cytokine receptors in the system are conserved through all simulations, see main text for details. **(C-D)** Sensitivity analysis of all key parameters with respect to the surface concentration average, gradient and surface concentration s.d., analogous to Figure 1F. **(E)** Correlation of fraction of secreting cells and signaling mode with the fraction of pSTAT+ cells. **(F)** Gradient, surface concentration s.d. and fractions of pSTAT5+ cells over the percentage of all receptors on responding cells. Low percentages correspond to autocrine, high to paracrine signaling.

##### Figure S3, related to Figure 2 in the main text.

**(A-B)** surface concentration s.d. and surface concentration average over (A) the fraction of secreting cells and (B) receptor heterogeneity, analogous to Figure 2B. Additionally a comparison of average concentration over receptor heterogeneity of the RD- and well-mixed system with the analytical approximation of (28) is shown. **(C)** Surface concentration average over the secretion rate of secreting cells for saturated and linear uptake functions. **(D-E)** Sensitivity analysis of the parameters under study with respect to surface concentration s.d., gradient and fraction of pSTAT+ cells, analogous to Figure 2C.

##### Figure S4, related to Figure 3 in the main text.

**(A)** Schematic of the delayed feedback process. To account for delays due to regulation or receptor expression auxiliary states are introduced (cf. Methods section in the main text). **(B-G)** Kinetic simulations, analogous to Figure 3B. Shown are (B) the % pSTAT5+ cells over the amount of secreting cells and varying feedback fold change $\gamma$ and (C-G) surface concentration average, receptors and pSTAT over time. Thin lines indicate the respective value of each cell in the system, bold lines the mean. **(H-I)** Surface concentration s.d. over time and over the amount of secreting cells.

##### Figure S5, related to Figure 4 and 5 in the main text.

**(A)** Cartoon visualization of the quantities niche score and niche effect, corresponding to Figure 4D. **(B-C)** Correlation between niche score and niche effect for varying fractions of secreting cells. The black line shows a linear fit of all data points. The arrow head was added to indicate the direction of increased feedback fold change. rs is the computed Spearman's rank correlation coefficient. **(D)** Silhouette score over clustering strength $\varphi$. **(E-G)** Surface concentration average and signaling range over the clustering strength of secreting cells for the systems under study in Figure 5.

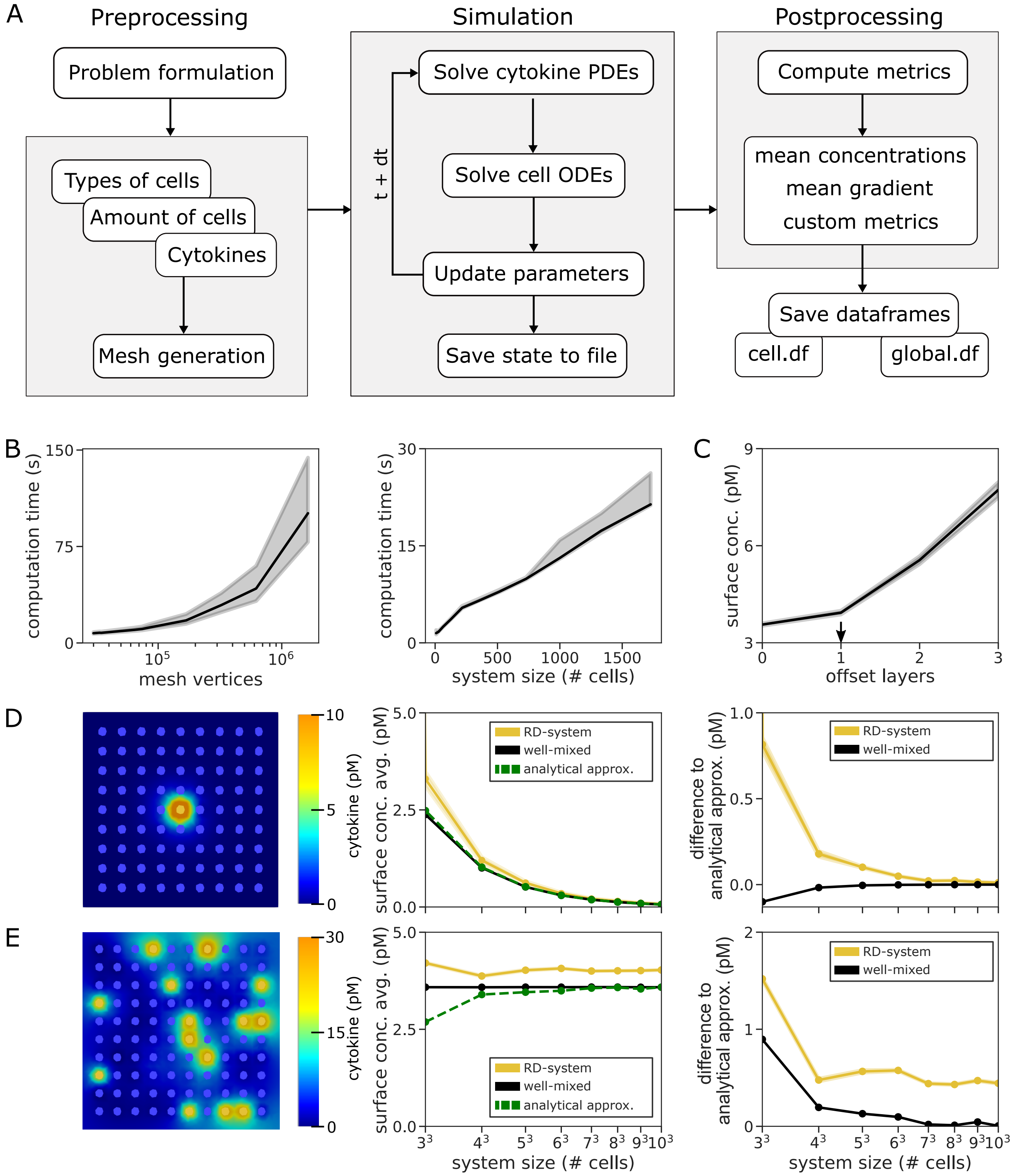


Brunner et al. Figure S1

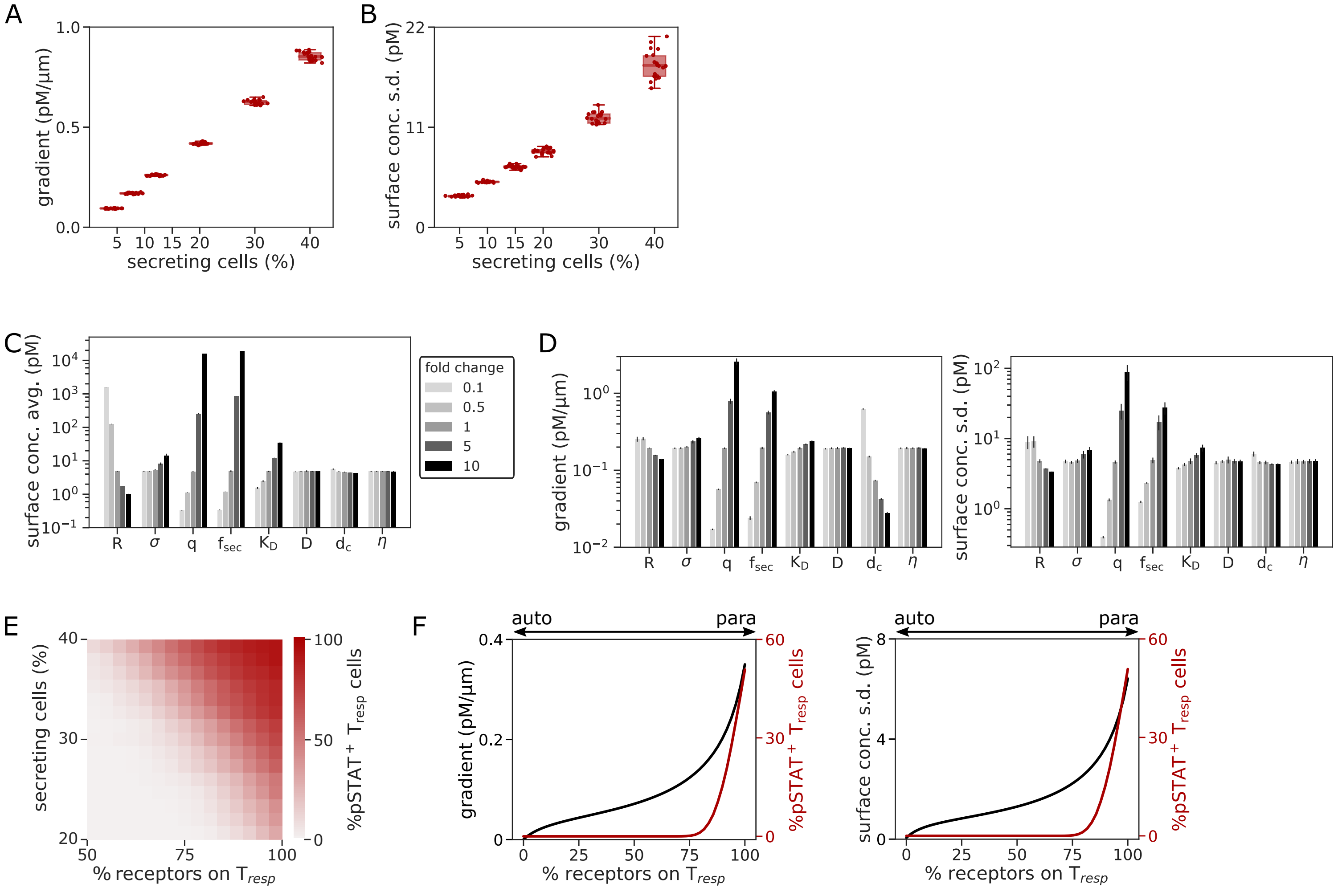


Brunner et al. Figure S2

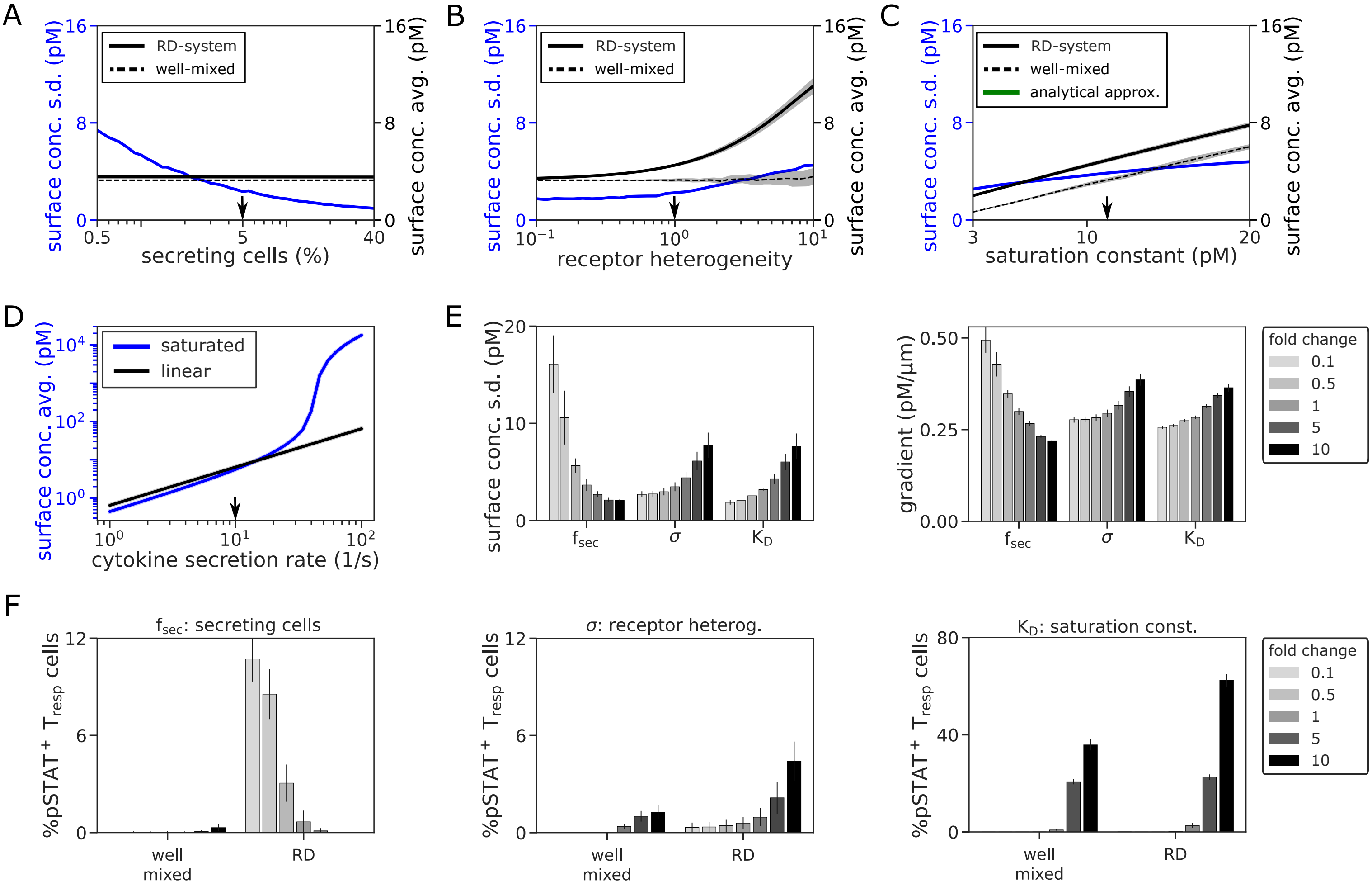


Brunner et al. Figure S3

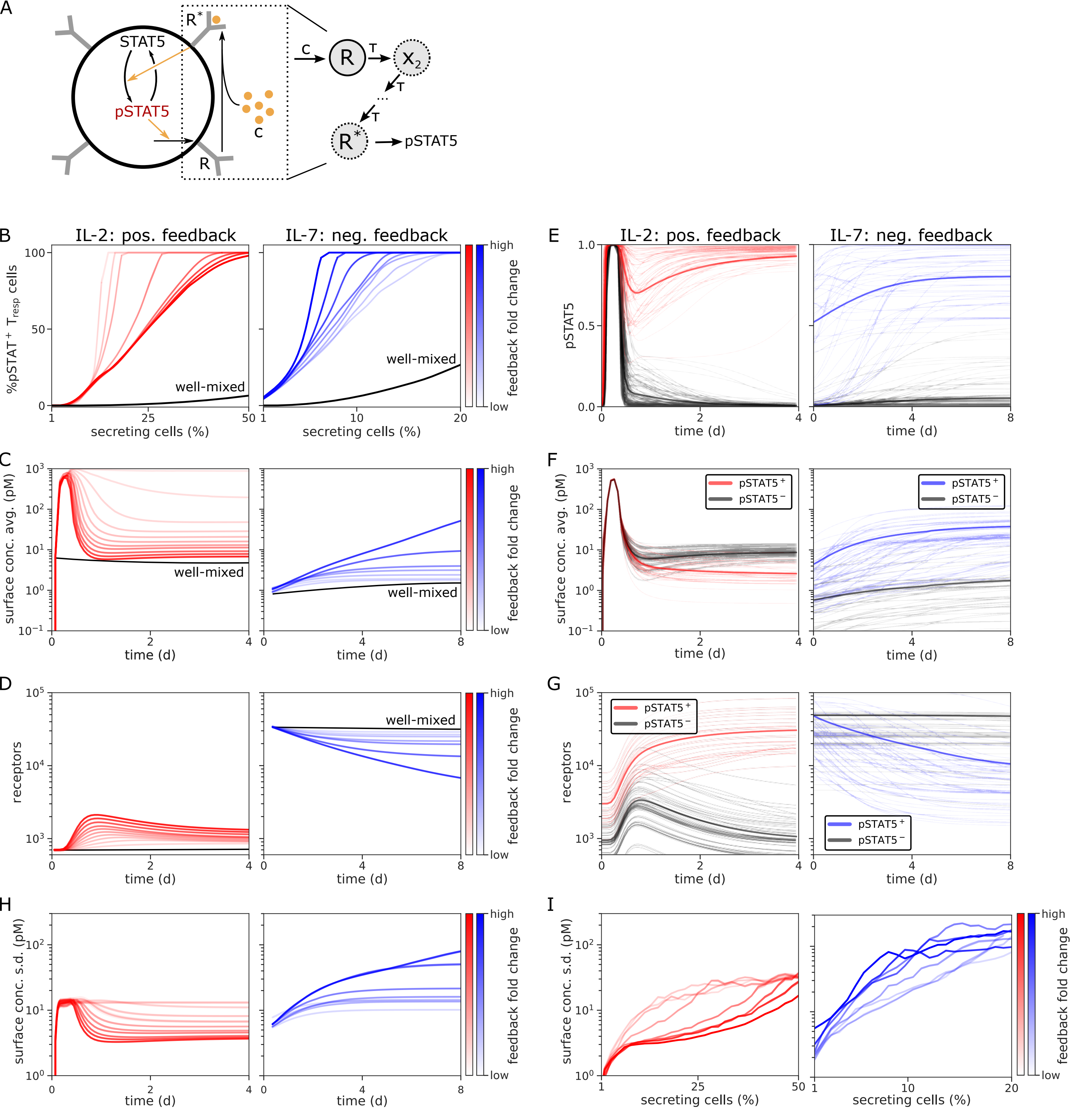


Brunner et al. Figure S4

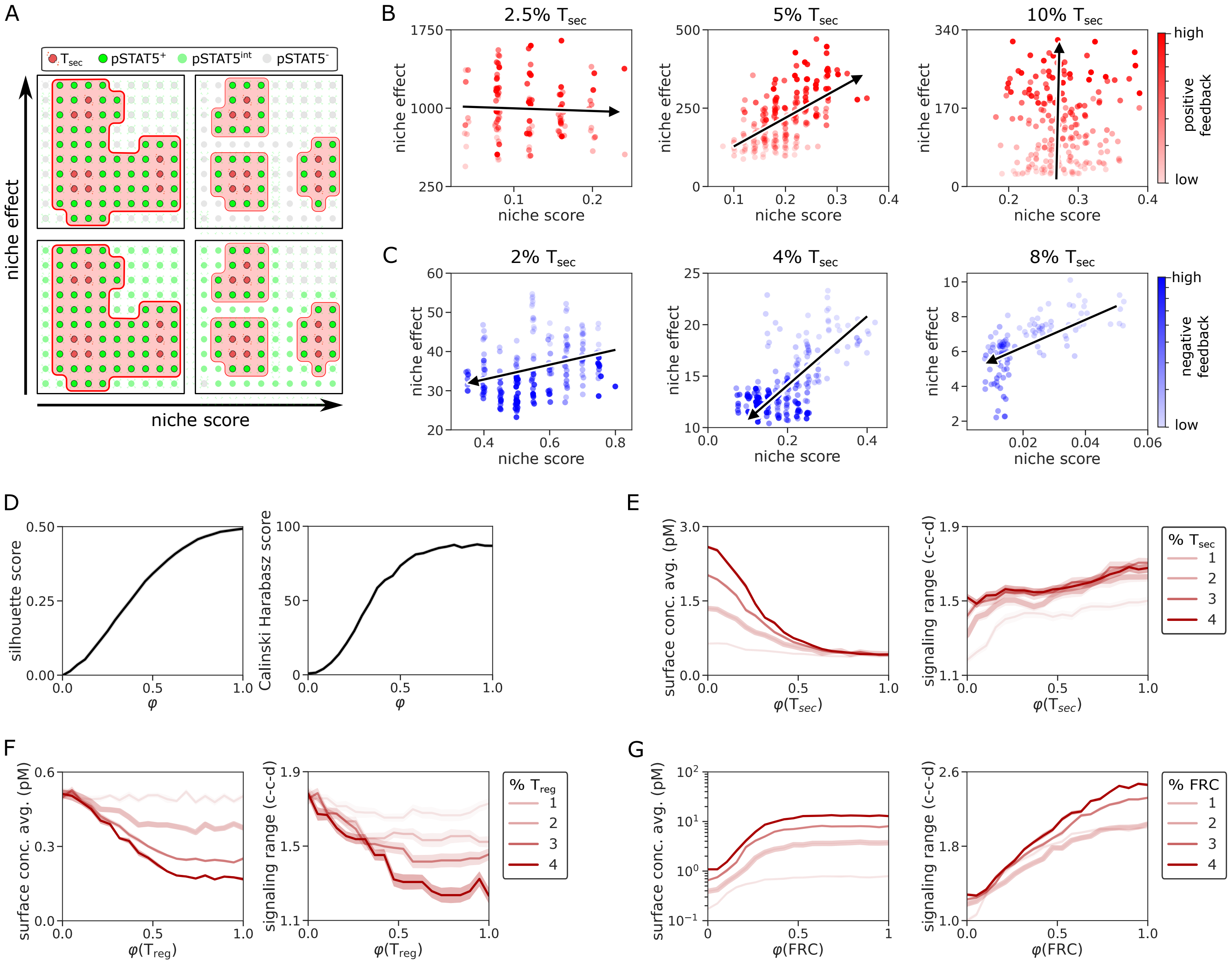


Brunner et al. Figure S5
